# Metagenomic Insights into The Diversity and Functions of Microbial Assemblages in Tasik Kenyir Ecosystem

**DOI:** 10.1101/176453

**Authors:** Mohd Ezhar Mohd Noor, Sharifah Noor Emilia Syed Jamil Fadaak, Mohd Noor Mat Isa, Mohd Faizal Abu Bakar, Muhd Danish-Daniel Abdullah

## Abstract

Tropical freshwater lake such as Tasik Kenyir are underrepresented among the growing number of environmental metagenomic data sets. In Tasik Kenyir, water from two different sites, pristine and disturbed areas were sampled. After the filtration process, genomic DNA from both sites were extracted using Meta-G-nome DNA isolation kit and shotgun metagenomic sequencing was carried out on Illumina HiSeq2500 Desktop Sequencer (Illumina, Inc.). Raw data were then trimmed and assembled using Metagenomic Assembler program, MetaVelvet. Data analysis was carried out using software Blast2GO (BioBam Bioinformatic S.L). The total number of sequence reads was 189,158 from TKS1.5m (disturbed area) and 246,577 from TKS2.5m (pristine area).The results indicate that sequence reads of microbial species were presence at disturbed area near the aquaculture zone was lower than the sequence reads of microbial species were presence at pristine area. When compared to archaea, both samples were dominated by bacteria (more than 90%) suggesting that bacteria are absolutely dominant in the prokaryotic communities in the freshwater samples. The lake appears to contain a mixture of autotrophs and heterotrophs capable of performing main biogeochemical cycles like nitrogen fixation by *Klebsiella* sp for TKS1.5m and *Pontibacter* sp. for TKS2.5m. and carbon fixation by heterotrophic *Alcaligenes* sp. and *Shewanella decolorationi* in TKS1.5m, and by *Pantoea* sp. in TKS2.5m. Present study will advance our understanding of the importance of freshwater microbial communities for ecosystem and human health.

## Introduction

All biogeochemical cycles on biosphere involve with the microbes and major redox reactions such as the carbon cycle is catalyzed by a set of key microbial enzymes (Falkowski *et al*., 2008). Microbes generally can be found everywhere in the biosphere, including soil, water bodies and sediments. Szabo-Taylor *et al.,* (2010) stated that microbes are widely known for their part in controlling the ecosystem function and biogeochemical cycling in the biosphere. It has been proposed by Newman and Banfield, (2002) that naturally, most microbes live under either nutrient-or energy-limiting conditions and due to this, they need to exist and blend in complex communities with the other microorganisms within the environments in order to survive.

Freshwater bodies such as Tasik Kenyir provide various ecological habitats and environment resources to flora and fauna. According to Portal of Lake Kenyir, (2014), Tasik Kenyir is the largest man-made lake in South East Asia and holds around 23.6 million cubic meters of water and occupies about 38000 hectares and also serves as another gateway to Malaysia National Park. Previous study by Amann *et al*., (1995) suggested that approximately less than 1% of earth’s microorganisms are cultivable. In addition, Rinke *et al*., (2013) also reported the majority of the cultivated and genome-sequenced bacteria belong to only four phyla (*Proteobacteria*, *Firmicutes*, *Actinobacteria*, and *Bacteroidetes*). Meanwhile, recent data had suggested the existence of at least 63 bacterial phyla and candidate groups (SILVA database version 115) as reported by Pruesse *et al.,* (2007) and Quast *et al*., (2013). These studies has given the researcher a better insight on the undiscovered genetic potential concealed by bacteria and archaea that evolved in billions of years. Therefore, a metagenomic analysis was done in order to compare the community compositions in Kenyir Lake as well as their role and functions.

Metagenomics is the technique used to extract the total genomic DNA of microorganisms directly from the environment and will represents the vast majority of microorganisms on earth as stated by Hugenholtz *et al*., (1998). The new generation of sequencing technology, with its ability to sequence thousands of organisms in parallel, has proved to be uniquely suited to this application. Metagenomics can be considered a revolutionary approach to study the microbial community that is unapproachable by available conventional methods and this approach also can capture the total genomes that present in a community of interest. As stated by Schloss and Handelsman, (2003), metagenomic was builds on advances in microbial genomics and in the polymerase chain reaction (PCR) amplification and cloning of genes. The field of metagenomic has play a major roles for in significant progress onmicrobial ecology, evolution, and diversity over the past 5 to 10 years. This development has opened new ways of understanding microbial diversity and functions. Hence, the characterization of the microbial community of Tasik Kenyir has potential implications for human activity and health which may apply to other freshwater lakes.

## Materials and Methods

### Water Sampling

Water samples were collected on January 2015 from Sungai Lasir (pristine area; N04°59.997 E102°50.133) Sungai Como (disturbed area; N05°01.887 E102°50.600) i.e a transect between Mentong River and Terenggan River and another transect between Como River Aquaculture Zone and Sultan Mahmud Hydroelectricity Power Plant. Water from these areas were collected at the surface level, approximately 5 meters from the water surface.

About 500 mL to 10 L of water were filtered through a filter cartridge. A membrane filter tower was constructed from polycarbonate filtration units (Millipore) with a 0.22 μm pore size membrane filter (Millipore) aseptically placed on the filter holder. Membrane filters were sterilized by autoclaving. After filtration, samples were snap-frozen immediately in liquid nitrogen and stored in -80°C until further use.

### Metagenomic Analysis

Nucleic acid extractions were performed by using Metagenome DNA isolation kit (Epicentre) and PowerWater DNA Isolation Kit (MO BIO Lab. Inc.). After that, gel Electrophoresis and Documentation was run to check the Quality and Quantity of the DNA. Shotgun metagenomic sequencing was carried out on Illumina HiSeq2500 Desktop Sequencer (Illumina, Inc.). The sequencing was performed by Macrogen Inc., Korea. Illumina platform was chosen due to lower cost and limited systematic errors compared to other platform. The generated multi-million reads were then trimmed and assembled by Metagenomic Assembler program, MetaVelvet. Data analysis was carried out using software Blast2GO (BioBam Bioinformatic S.L). Then initial taxonomic analysis was performed by bioinformatic tools like BLAST (Altschul *et al*. 1997) and MEGAN (Huson *et al*. 2007).

## Results and Discussion

Sequence reads data from two sites were collectively used to analyze microbial diversity. The total number of reads was 189,158 from TKS1.5m (disturbed area) and 246,577 from TKS2.5m (pristine area) (Table 1). When compared to Archaea, both samples were dominated by Bacteria with more than 90%, suggesting that bacteria are absolutely dominant in the prokaryotic communities in the freshwater samples. Phylogenetic analysis of both samples also revealed that the overwhelming majority of the data set were bacterial and archaeal. Most of the archaeal metagenomics reads from TKS1.5m area matched *Methanosaeta thermophile* with a few additional representatives within Thermococci while archaeal metagenomics reads from TKS2.5m area matched *Methanolobus psychrophilus.* Both metagenomics data sets agreed with the broad taxonomic picture that almost 90% of the sequences affiliated to Bacteria, whereas Archaea were a minor component. The remaining about 10% sequences were of eukaryotic origin and virus. All major bacterial groups commonly found in freshwater ecosystems that has been reported by Humbert *et al.,* (2009) were present in Tasik Kenyir.

**Table 1.** Characteristics of shotgun metagenomic libraries

| Characteristic | Sample designation |  |
| --- | --- | --- |
|  | TKS1.5m | TKS2.5m |
| Total Reads | 189158 | 246577 |
| Genes Reads | 66338 | 106097 |
| Species Detected | 4153 | 32421 |

Shotgun metagenomic analysis revealed that both samples were dominated by Proteobacteria (more than 65%, Table 2). The finding is consistent with findings of past studies by Kersters et al., (2006), which reported that Proteobacteria, the largest and most phenotypically diverse phylum, accounting for at least 40% of all known genera. In addition, present study, phyla in common to all TKS1.5m and TKS 2.5m samples (Verrucomicrobia, Bacteroidetes, and Firmicutes, Cyanobacteria; Table 2) can be found in a variety of aquatic environments similar to the finding by Dillon *et al*., (2009) although latter phyla were far less dominant than the Proteobacteria. In contrast to sample fom pristine area, TKS2.5m, sample from disturbed area (aquaculture site), TKS1.5m was dominated by pathogenic bacteria species like Pseudomonas species, Aeromonas species, Klebsiella species and Vibrio species (Figure 3). Austin and Austin (1999) stated that common fish pathogenic bacterial species belong to the genera *Vibrio*, *Aeromonas, Flavobacterium, Yersinia, Edwardsiella* and *Pseudomonas*. In addtion, study by Martin-Carnahan and Joseph (2005) also stated that Aeromonads are among the most common bacteria in aquatic environments and are frequently associated with severe diseases among cultured fishes.

**Table 2.**
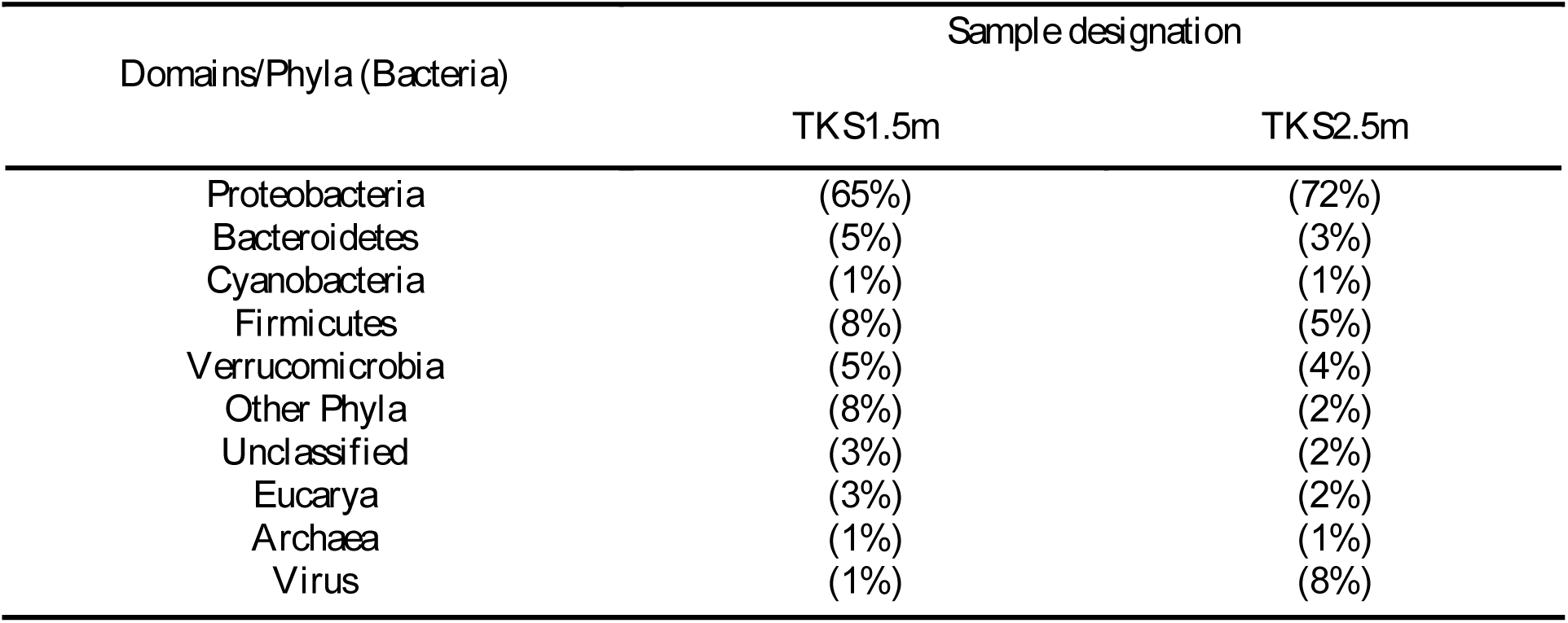
Percentage of Domains and bacterial phyla from TKS1.5m and TKS2.5m

**Figure 1:**
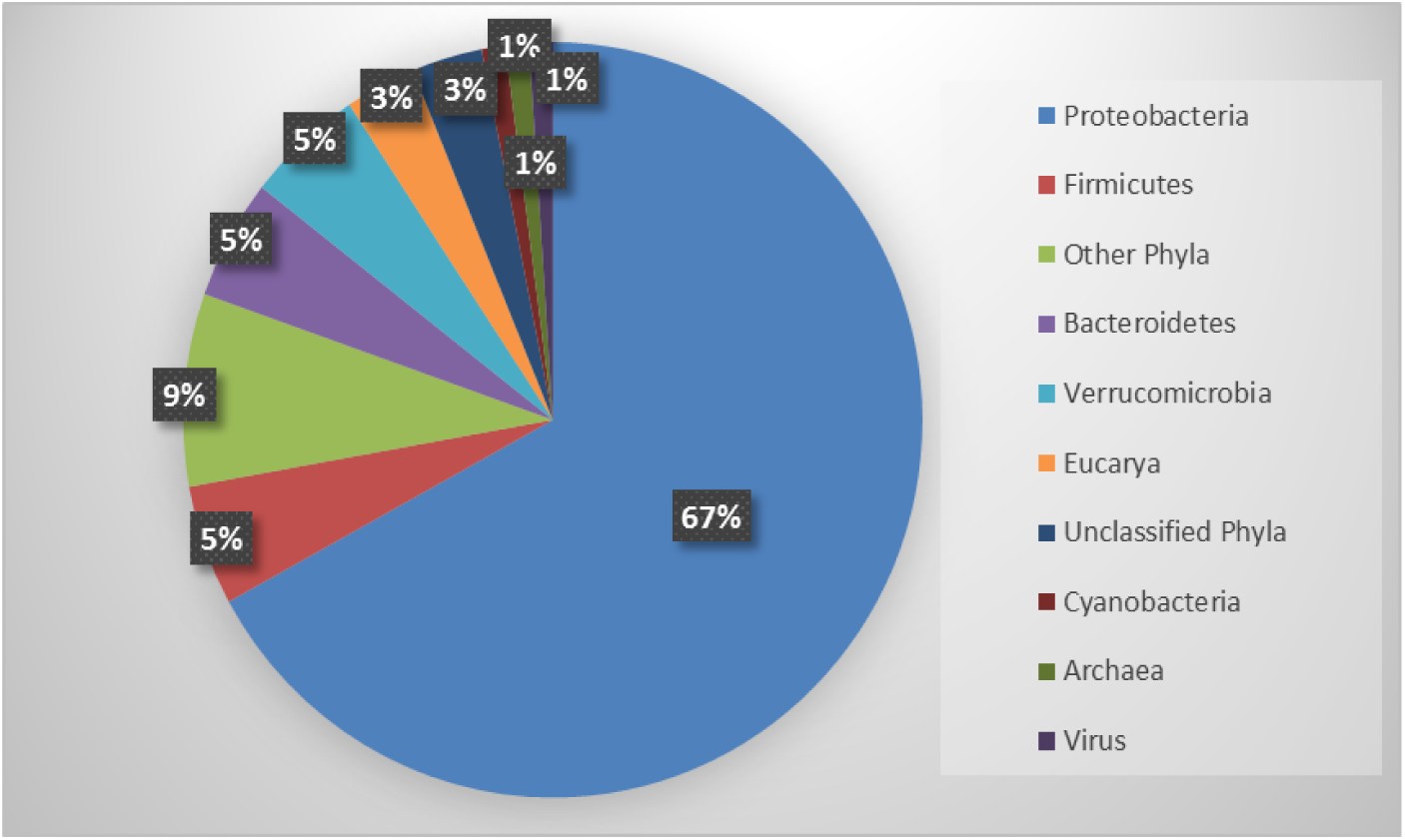
Relative abundance of the domain and dominant bacterial phyla from TKS1.5m

**Figure 2:**
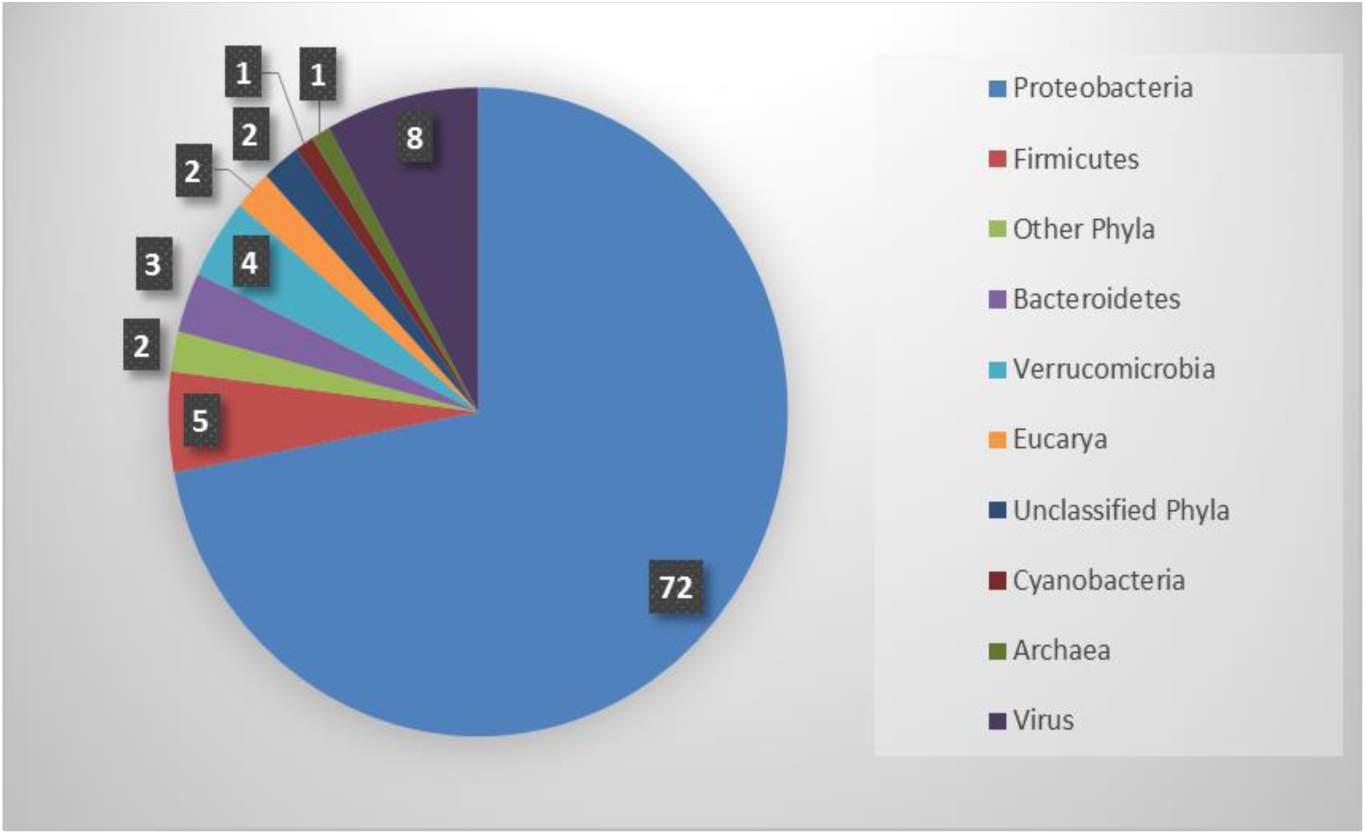
Relative abundance of the domain and dominant bacterial phyla from TKS2.5m.

**Figure 3:**
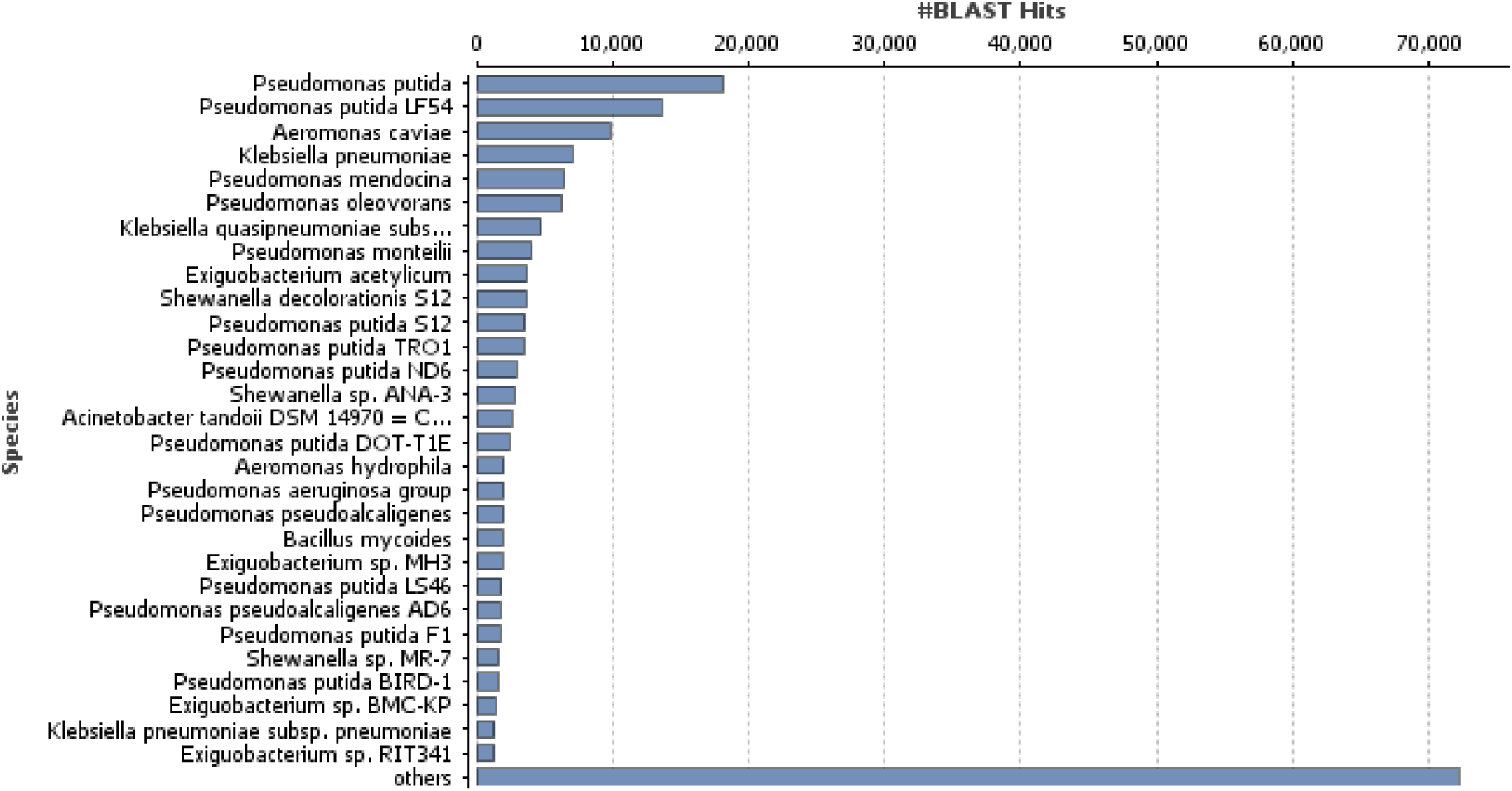
Species Distribution of microbial community detected from the TKS1.5m (Aquaculture Area) metagenome sequences.

**Figure 4:**
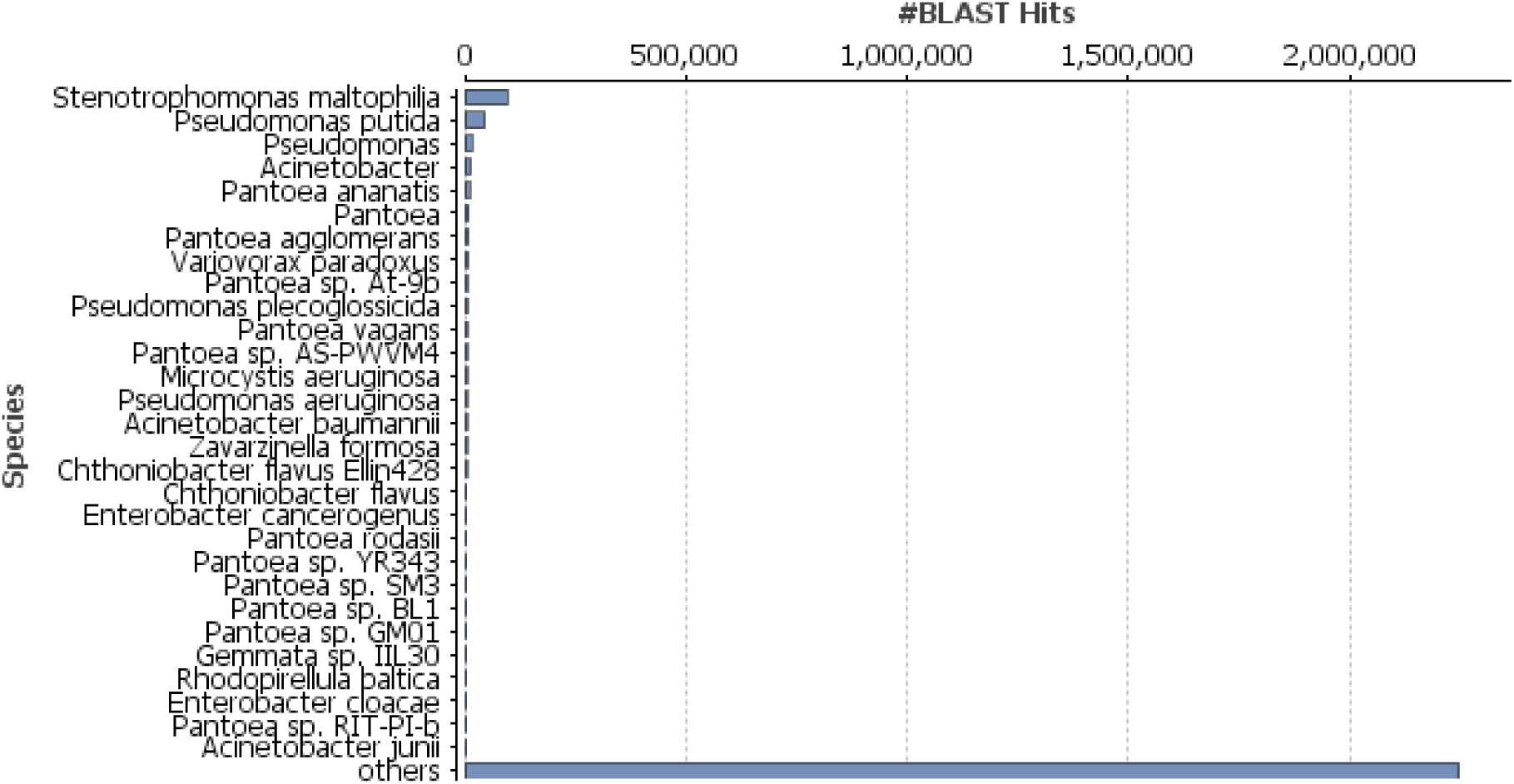
Species Distribution of microbial community detected from the TKS2.5m (Pristine Area) metagenome sequences.

As known, Bacteria in the genus *Vibrio* are mainly pathogenic to marine and brakishwater fish. However, the scientific report by Lightner and Redman (1998) stated they are occasionally reported in freshwater species. Research finding by Actis *et al.,* (2011) also points toward that vibriosis, one of the major bacterial diseases affecting fish is primarily caused by pathogenic species such as *Vibrio harveyii, V. anguillarum* and *V. ordalii.* These vibrios can be found abundantly in the sample from aquaculture site, TKS1.5m compared to pristine area sample, TKS2.5m. Although pathogenic species representing majority of existing bacterial taxa, only a relatively small number of pathogens are responsible for important economic losses in cultured fish worldwide. Present result analysis also revealed that microbial community from TKS2.5m was more diverse than the TKS1.5m. Consistent with findings by Kennedy (1999), the diversity of microbial is critical to the functioning of the ecosystem, because there is the need to maintain ecological processes like controlling pathogens within the ecosystems. Study by Yamanaka *et al*. (2003) also stated that microbial diversity is directly related to ecosystem stability.

As Bacteria and Archaea accounted for most of total metagenomic reads, the comparative study of the geochemistry of carbon (C) and nitrogen (N) will be focused on the prokaryotes. For the carbon cycling, the pathway detected in the oxic–anoxic interface was aerobic respiration by heterotrophic *Alcaligenes* sp. from the order *Burkholderiales* and *Shewanella decolorationi* from *Alteromonadales* in TKS1.5m, and by *Pantoea* sp. from *Enterobacteriales* in TKS2.5m. Carbon cycling is a key metabolic feature of members of the alpha and gamma Proteobacteria, including *Alteromonadaceae*, *Pseudomonadaceae*, *Sphingomonadaceae*, and *Vibrionaceae* as well as *Flavobacteriaceae*, which is member of the Cytophaga-Flavobacteria-Bacteroides (CFB) clade. The present finding also support the previous studies by Ramaiah *et al.,* (2000), Ivanova and Mikhailov, (2001), Kirchman, (2002) Kirchman *et al*., (2004) and Mikhailov *et al*., (2006) which concluded that these bacteria produce enzymes capable of degrading various polysaccharides, such as cellulose, chitin, and pectin. For the nitrogen cycle, most of the detected marker genes catalyzed Nitrogen fixation by *Klebsiella* sp for TKS1.5m and *Pontibacter* sp. for TKS2.5m. The result is in the lines of earlier literatures by Jenni *et al*., (1989), Willems *et al*., (1991) and Chimetto *et al*., (2008) which found that nitrogen fixation has been reported among members of the *Comamonadaceae*, *Pseudomonadaceae*, and *Vibrionaceae* families. Meanwhile, denitrification was observed in relatively low abundance in both samples but, metagenomic reads matching *Nitrospirae* nitrite-oxidizing bacteria were detected in both samples, pointing out that the genetic potential to close the nitrogen cycle was there. Studies by Paerl *et al*., (2002) and Rabalais *et al*., (2009) stated denitrification is important in aquatic environments because a build-up of nitrate can lead to eutrophication and harmful algal blooms. Earlier studies by Cole, (1996), Chou *et. al*., (2008) and Rajakumar *et al*., (2008) showed that nitrogen cycling is the key metabolic trait shared by different bacteria associated with both samples where many of the bacteria, including members of the *Comamonadaceae*, *Enterobacteriaceae* and *Pseudomonadaceae*, can reduce nitrate.

## Conclusion

It is known in present study that prokaryote community bacteria is absolutely dominant over archaea with more than 90% in metagenomic pools in the freshwater prokaryotic samples. Tasik Kenyir metagenomic data sets provides the means to examine with fine-scale resolution how natural microbial populations are impacted by human activities (aquaculture site) and major environmental perturbations. Hence, it enable researchers to identify some of the major microbial players and functions that could be targets for future work examining the response and resilience of these communities to environmental changes and human perturbations. Such studies will significantly advance the understanding of the importance of microbial communities for ecosystem and human health especially in freshwater environment.

## Acknowledgements

This research was funded by the Ministry of Higher Education Malaysia through grant FRGS 2014 Vot No. 59334. We thanks all the people involved directly or indirectly in this study especially staffs from the School of Fisheries and Aquaculture Sciences, UMT, Mr Sharol Ali and senior scientist from Malaysia Genome Institute, Mr. Faizal Abu Bakar.

